# Clustered synapses develop in distinct dendritic domains in visual cortex before eye opening

**DOI:** 10.1101/2023.03.02.530772

**Authors:** Alexandra H. Leighton, Juliette E. Cheyne, Christian Lohmann

**Affiliations:** Department of Synapse and Network Development, Netherlands Institute for Neuroscience, 1105 BA Amsterdam, the Netherlands; Department of Functional Genomics, Center for Neurogenomics and Cognitive Research, VU University Amsterdam, the Netherlands

**Author notes:** Present address: Physiology Department, Centre for Brain Research, University of Auckland, Auckland, New Zealand (Juliette E. Cheyne).

## Abstract

Synaptic inputs to cortical neurons are highly structured in adult sensory systems, such that neighboring synapses along dendrites are activated by similar stimuli. This organization of synaptic inputs, called synaptic clustering, is required for high-fidelity signal processing, and clustered synapses can already be observed before eye opening. However, how clustered inputs emerge during development is unknown. Here, we employed concurrent *in vivo* whole-cell patch clamp and dendritic calcium imaging to map spontaneous synaptic inputs to dendrites of layer 2/3 neurons in the mouse primary visual cortex during the second postnatal week until eye opening. We find that the number of functional synapses and the frequency of transmission events increase several fold during this developmental period. At the beginning of the second postnatal week, synapses assemble specifically in confined dendritic segments, whereas other segments are devoid of synapses. By the end of the second postnatal week, just before eye-opening, dendrites are almost entirely covered by domains of co-active synapses. Finally, co-activity with their neighbor synapses correlates with synaptic stabilization and potentiation. Thus, clustered synapses form in distinct functional domains presumably to equip dendrites with computational modules for high-capacity sensory processing when the eyes open.

## Introduction

Building a functional brain requires neuronal circuits to be wired up with high specificity via synapses between connecting neurons. Developing neurons select their synaptic partners based on molecular cues such as adhesion molecules and subsequently refine their connections through activity-dependent synaptic plasticity, stabilization and elimination driven by spontaneous network activity and later, after the onset sensory function, by experience (Cline, 2003; Martini et al., 2021; Sanes and Zipursky, 2020). Electron microscopy revealed that excitatory synapses between neurons emerge during the end of the first postnatal week in mouse and rat sensory cortex (Blue and Parnavelas, 1983; De Felipe J. et al., 1997; Miller and Peters, 1981; Wildenberg et al., 2023). The addition of new synapses peaks at the end of the second postnatal week and synapse density in the primary visual and somatosensory cortices approaches adult levels once the eyes open at postnatal day (P) 14 (Blue and Parnavelas, 1983; De Felipe J. et al., 1997). While these structural studies clearly showed the importance of this developmental period for adding large numbers of synaptic connections in emerging circuits, how cortical dendrites establish functional inputs has not been investigated on the level of individual synapses in vivo.

In the adult cortex, synaptic inputs to dendrites are highly organized with sub-cellular specificity. Recent studies demonstrated that synaptic inputs of cortical pyramidal cells are clustered, such that synapses with similar activity patterns or stimulus selectivity are located near each other along their dendrites (Iacaruso et al., 2017; Ju et al., 2020; Kerlin et al., 2019; Otor et al., 2022; Scholl et al., 2021; Takahashi et al., 2012; Wilson et al., 2016; Winnubst et al., 2015). Since pyramidal cell dendrites integrate local inputs supra-linearly (Branco and Hausser, 2011; Harnett et al., 2012; Losonczy and Magee, 2006; Makara and Magee, 2013), this arrangement of clustered inputs is thought to allow for local synaptic integration in dendritic computational subunits (Larkum, 2022; Larkum and Nevian, 2008; Major et al., 2013; Tran-Van-Minh et al., 2015). Local dendritic integration dramatically increases the computational power of neurons (Poirazi and Mel, 2001). Furthermore, experimentally blocking supra-linear dendritic integration disturbs sensory stimulus sensitivity and selectivity in cortical neurons (Lavzin et al., 2012; Palmer et al., 2014; Smith et al., 2013; Xu et al., 2012). Finally, learning is associated with structural plasticity of clustered synapses (Cichon and Gan, 2015; Fu et al., 2012; Hedrick et al., 2022; Makino and Malinow, 2011; McBride et al., 2008). Together, these studies highlight the fundamental importance of the subcellular organization of synaptic inputs along dendrites.

This raises the question of how the precise subcellular organization of synapses is achieved during development. Synaptic clustering can be observed early on in the developing visual system of tadpoles and mice (Podgorski et al., 2021; Winnubst et al., 2015), suggesting that synaptic inputs develop with some degree of specificity, but the trajectory of functional synaptic input development has been unclear. Specifically, it is unknown (1) when synaptic inputs become clustered, (2) whether synapses are formed in a clustered state or arise in a random state to become sorted into clusters later, and (3) whether synapses with similar input patterns distribute smoothly along dendrites or emerge in distinct domains.

Here, we performed *in vivo* whole-cell patch clamp recordings with two-photon calcium imaging (Chen et al., 2011; Helmchen et al., 1999; Jia et al., 2010; Svoboda et al., 1997; Takahashi et al., 2012; Winnubst et al., 2015) of mouse primary visual cortex layer 2/3 neurons, allowing us to map functional synaptic inputs across dendritic arborizations in the course of the second postnatal week. We combined resonant scanning with piezo-driven focusing to image large proportions of a neuron’s dendrites. The total dendritic length imaged per neuron was almost ten times larger than in our previous study (Winnubst et al., 2015), allowing us to map the large-scale distribution of synapses along developing dendrites.

We found that the number of functional synapses and the frequency of transmission events at individual synapses increase rapidly before eye opening. Furthermore, we observed that clustered synaptic inputs accumulate in spatially and functionally separated domains already at the beginning of the second postnatal week. By the end of the second postnatal week, dendrites had become almost entirely covered by functional domains of co-active neighbors, most likely through local plasticity driven by coincident spontaneous network activity. Thus, already before eye opening functional domains develop in cortical dendrites, which can serve as distinct computational modules for processing visual stimuli thereafter.

## Results

### Mapping synaptic inputs onto spine and shaft synapses

To visualize the activity and spatial organization of functional excitatory synapses in the developing mouse primary visual cortex, we imaged spontaneously occurring synaptic calcium transients in apical dendrites of pyramidal cells in L2/3 using the calcium indicator GCaMP6s in neonatal mice (Fig 1A). We combined resonant scanning and piezo-driven z-positioning to image a large area of the dendritic tree of individual neurons (Fig 1B). Calcium transients at individual synapses reflecting synaptic transmission (Jia et al., 2010; Takahashi et al., 2012; Winnubst et al., 2015) allowed us to map synaptic inputs across 12 dendritic areas from 11 mice at ages between postnatal day 8 to 13, the day before eye opening. Together, these dendrites hosted 354 functional synapses which produced 3440 synaptic transmission events. We recorded these neurons in voltage clamp configuration to measure barrages of synaptic currents arriving at the soma simultaneously with individual synaptic transmission events monitored by calcium imaging (Fig 1C).

**Fig. 1:**
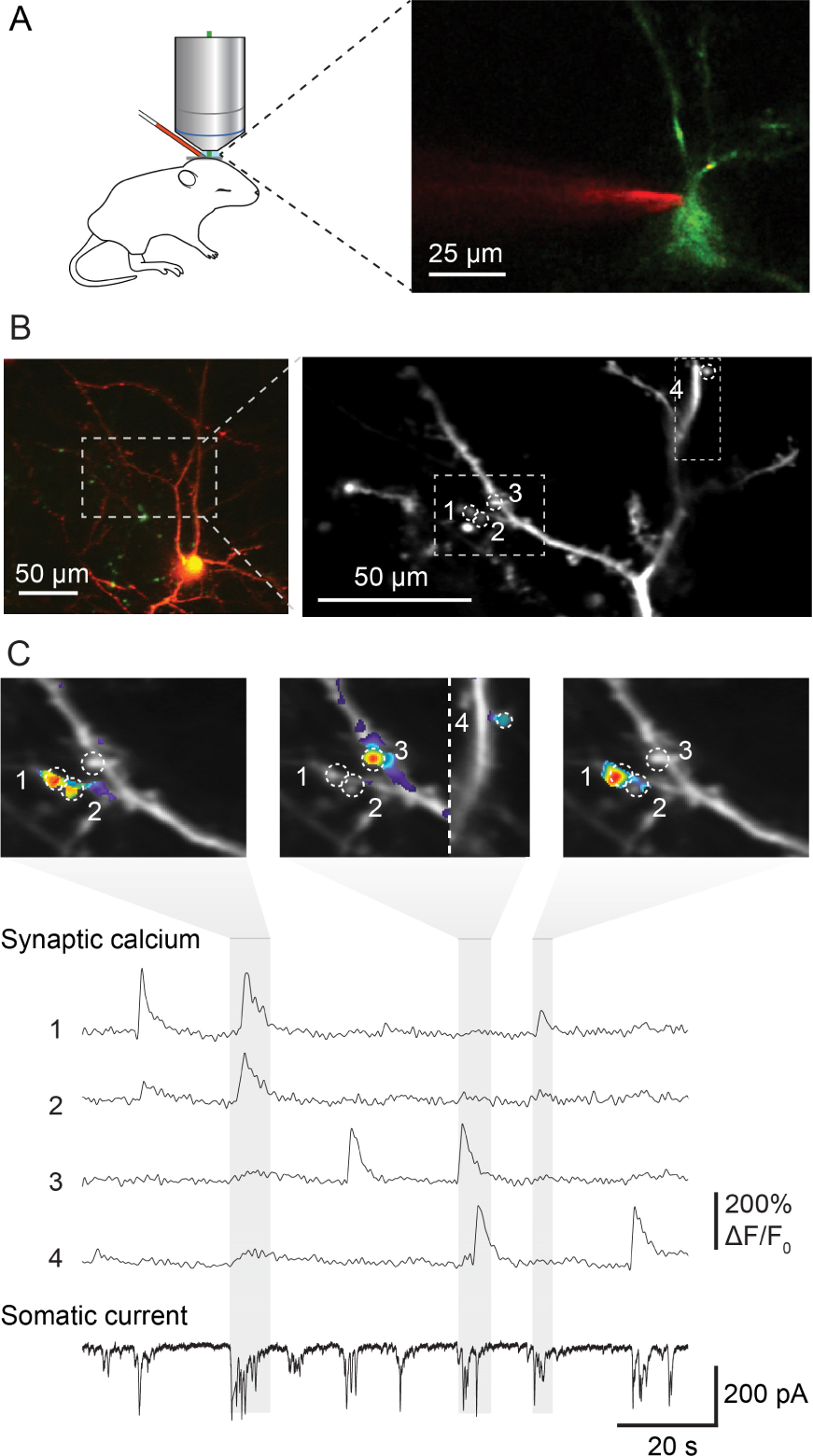
Mapping functional synaptic inputs of V1 layer 2/3 pyramidal cell dendrites in vivo. A. Schematic of experimental setup (left) and pyramidal cell in mouse visual cortex expressing DsRed and GCaMP-6s (right). Layer 2/3 neurons were targeted in vivo under two-photon guidance. Patch-clamp pipettes were coated with Alexa 594 for visualization. B. Left: Layer 2/3 pyramidal neuron expressing DsRed and GCaMP-6s at postnatal day (P) 12 (D9 in Suppl. Figures 1 and 2). Right: High magnification view of the dendrite marked with dashed lines on the left. Dashed circles and numbers mark four individual synaptic sites. Dashed rectangle indicates views shown in C. C. Spontaneous synaptic inputs were visible both as local increases in fluorescence at the four synaptic sites labeled in B, and as synaptic currents in the somatic whole-cell voltage-clamp recording. Grey vertical bars indicate three individual bursts, and panels above show synaptic calcium increases at the four example synapses during each labeled barrage. Synaptic transmission of individual synapses could be detected and distinguished clearly from that at neighboring synapses.

In the mature cortex, most excitatory synapses of pyramidal neurons are located on spines (Berry and Nedivi, 2017; Kasai et al., 2021; Moyer and Zuo, 2018). Spines provide some chemical and electrical isolation from the dendrite’s shaft and produce clearly detectable calcium signals evoked by presynaptic inputs (Fig 2A). However, during development, many cortical synapses are formed directly onto the shaft (Miller and Peters, 1981; Wildenberg et al., 2023) where synaptic calcium transients are masked by calcium influx triggered by back-propagating action potentials. Therefore, we prevented action potential firing by recording layer 2/3 pyramidal neurons in voltage-clamp mode during imaging and the intracellular sodium channel blocker QX314 in the patch-pipette. The holding potential was set to –30 mV to facilitate calcium influx through NMDA receptors (Jia et al., 2010; Takahashi et al., 2012; Winnubst et al., 2015). In this configuration we could image local calcium transients at both spine and shaft synapses (Fig 2A, B, see Methods for details and potential caveats) and map all functional excitatory synaptic inputs onto these dendrites across space and time (Fig 2C, Supplementary Figures 1 and 2).

**Fig. 2:**
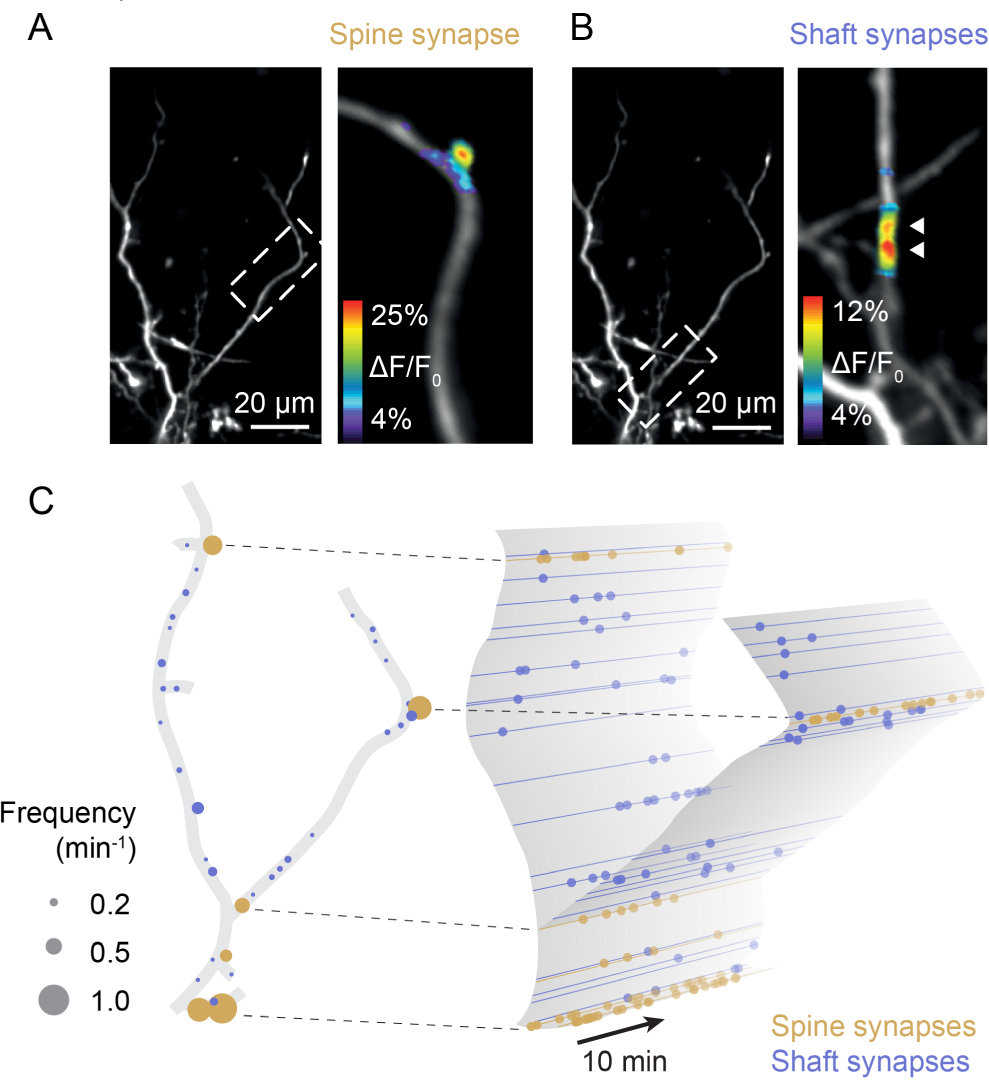
Sampling synaptic transmission in spine and shaft synapses (D3). A. Synaptic transmission in a spine of a layer 2/3 pyramidal neuron at P9. B. Synaptic transmission at two neighboring shaft synapses of the neuron shown in A. C. Schematic representation of the synaptic inputs of the neuron shown in A and B. Left: All functional synapses are labeled as individual discs. Yellow and blue discs represent spine and shaft synapses, respectively. The disc diameter indicates the observed transmission frequency at each synapse. Right: Schematic representation of synaptic transmission across time in the same dendrite. Dots represent individual synaptic transmission events.

### Synaptic inputs increase rapidly during the second postnatal week

Previous electron microscopy studies revealed that synapses form at the highest rate during the second postnatal week in rodent sensory cortex (Blue and Parnavelas, 1983; De Felipe J. et al., 1997; Wildenberg et al., 2023); however, the course of functional synapse development *in vivo* has not been investigated yet. In line with these previous EM studies, we report here that the density of functional synapses increased significantly between P8 and P13 (Fig 3A-B). The frequency of synaptic transmission at individual synapses increased as well (Fig 3C). Hence, the number of synaptic transmission events in a given dendrite was higher by P12/13 (n = 6 dendrites) compared to the beginning of the second postnatal week (P8-10; n = 6 dendrites; Fig 3D). While the percentage of spine synapses was slightly higher in older dendrites, there was no significant relationship between age and the percentage of spine synapses across the sampled dendrites (Fig 3E). This is in line with previous observations that spine development is maximal after P14 (Summarized in: Lohmann and Kessels, 2014).

**Fig. 3:**
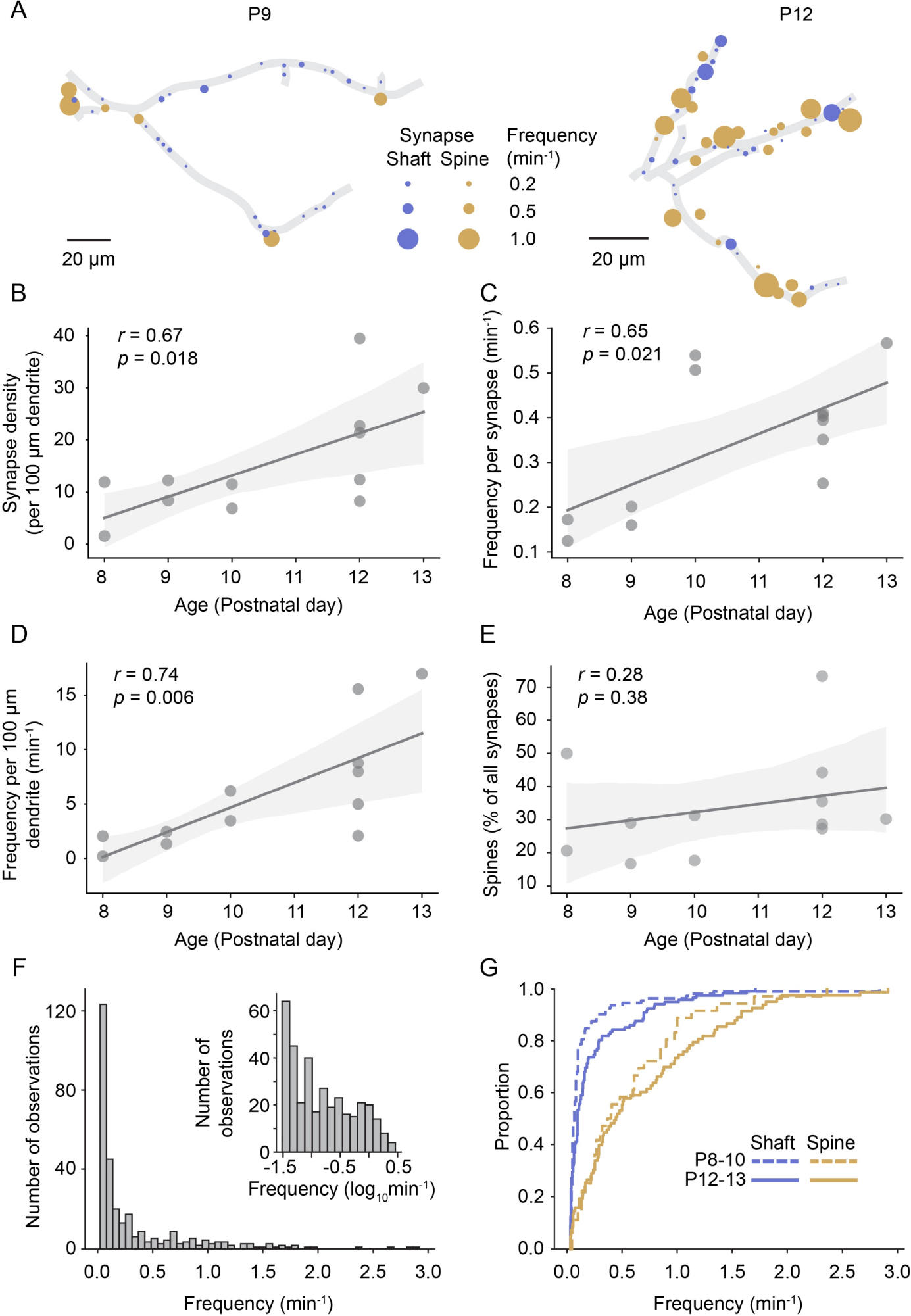
Synapse density and synaptic transmission frequency increased rapidly during the second postnatal week. A: Plots of layer 2/3 pyramidal cell dendrites from two experiments at P9 (D3) and P12 (D10), respectively. Each functional synapse is indicated as a disc where the diameter represents the frequency of transmission at shaft (blue) and spine (yellow) synapses. B: The density of functional synapses increased with age. Each dot represents the number of synapses per 100 micrometer dendrite of one experiment. Statistical parameters in this panel and C-E are derived from linear regressions on the data from n = 12 experiments. The light grey area indicates the 95% confidence interval as in all following regression plots. C: The frequency of synaptic transmission per synapse increased with age. Each dot represents the mean frequency across all synapses from one experiment. D: The frequency of synaptic inputs along a stretch of dendrite increased multi-fold during the second postnatal week. E: The percentage of synapses that are located on spines did not change significantly during the observed period. F: The distribution of transmission rates across all sampled synapses. While most synapses show only low transmission frequency, the distribution is very long-tailed, demonstrating that few synapses show very high synaptic transmission rates. Inset: frequency distribution shown on a logarithmic scale. G: Cumulative distribution of transmission frequencies across spine and shaft synapses in younger (P8-10) and older dendrites (P12-P13). The transmission frequency was higher at spine than at shaft synapses at both ages (each p < 10^-9^, Mann-Whitney-U tests, P8-10: n=36 spine / 113 shaft synapses, P12-13: n = 83 spine / 122 shaft synapses).

The distribution of synaptic transmission frequencies of all synapses showed that the majority of synapses received inputs at low frequencies, however, the long tail of the distribution demonstrated that a number of synapses were very active (Fig. 3F). In general, spine synapses were more active than shaft synapses in both young (P8-10; 0.55 ± 0.09 min^-1^ vs. 0.16 ± 0.03 min^-1^, mean ± SEM) and older animals (P12-13; 0.67 ± 0.07 min^-1^ vs. 0.23 ± 0.03 min^-1^; Fig 3G). While we detected synaptic transmission at the majority of spines, a fraction of structural spines remained inactive during our recordings in both younger (25 ± 20%) and older dendrites (9 ± 9% for recording durations of 29 ± 8 and 26 ± 6 min, respectively; mean ± STD), possibly representing presynaptically silent synapses (Voronin and Cherubini, 2004).

Next, we investigated when individual synapses were activated in relation to the barrages of spontaneous synaptic inputs neurons receive during this developmental period (Fig 4A). Barrages were identified as transient inward currents of at least 10 pA, which consisted of multiple synaptic currents. We found that most synaptic inputs occurred in barrages (59% in young neurons, 61% in older neurons). The rate of barrages did not change significantly with age (Fig 4B); however, we did observe a trend towards an increase in synaptic charge transferred per barrage (Fig 4C), as well as a significant increase in the number of synaptic transmission events per barrage (Fig 4D) during development. As expected, the number of events observed per barrage correlated highly linearly with the charge transferred during each barrage (Fig 4E). In addition, the amplitude of individual synaptic transmission events was higher during larger barrages (Fig 4F), suggesting that multiple synaptic vesicles were released simultaneously during larger barrages.

**Fig. 4:**
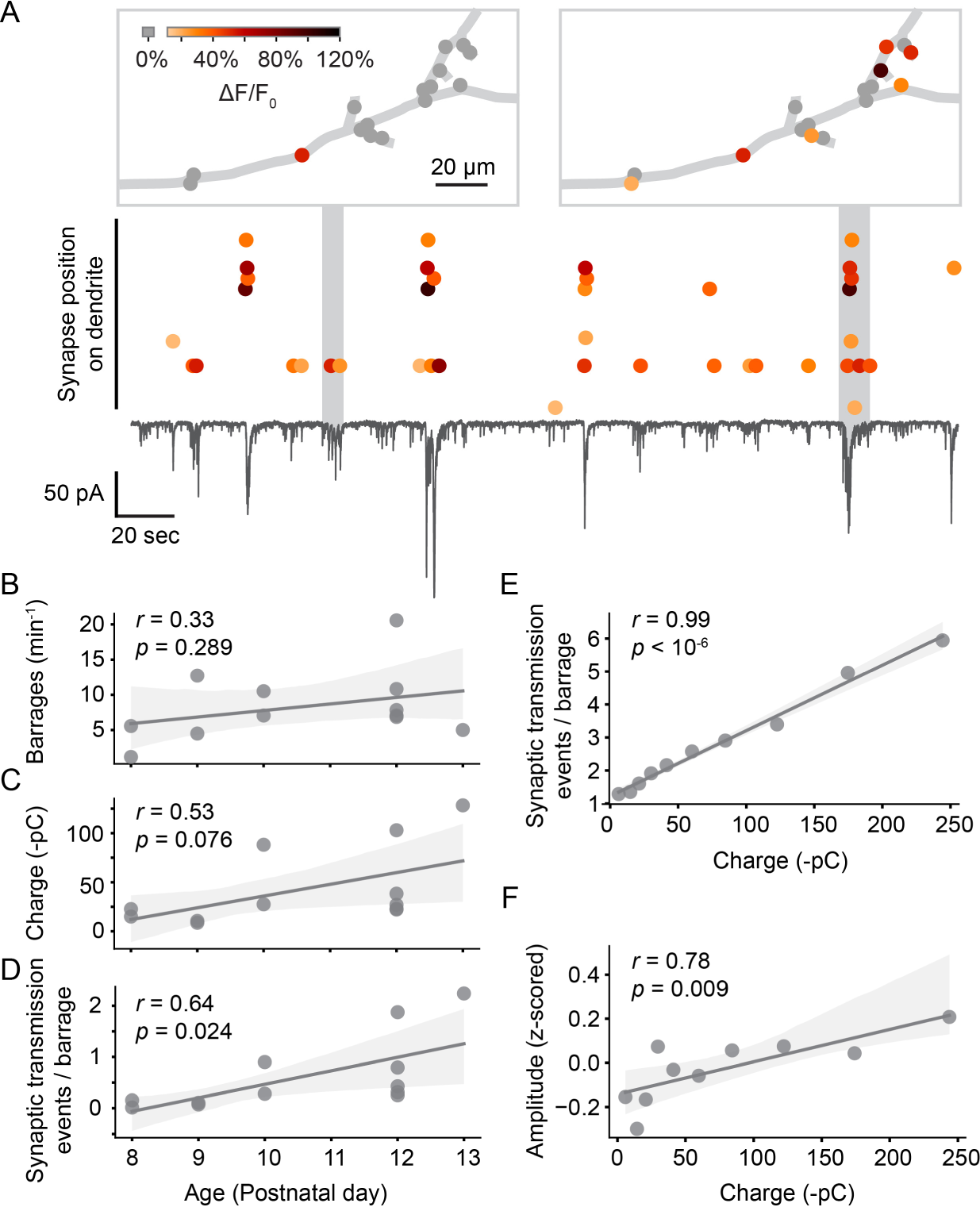
Synaptic transmission at the synapse and soma level. A: Synaptic transmission events (dots) at individual synapses and synaptic currents recorded simultaneously from the soma (black trace) in a layer 2/3 neuron at P10 (D6). Top: dendritic location of synapses that are active during two example barrages. Colors represent the amplitude of individual synaptic transmission events. Grey dots show synapses that were inactive during the respective barrages. Most synaptic transmission events occurred during barrages of somatic currents. B: The number of barrages did not change significantly during the observed developmental period. Each dot represents the mean from one experiment. The light grey area indicates the 95% confidence interval as in all following regression plots. C: The total charge transferred during barrages of synaptic currents showed a trend towards increases with age. D: The number of synaptic transmission events observed during individual barrages increased with age. E: The number of synaptic transmission events correlated linearly with the total charge transferred of barrages during which they occurred. Each data point shows the mean number of synaptic transmission events per barrage for geometrically binned barrage sizes. F: The amplitude of synaptic transmission events at a given synapse correlated with the total charge transferred, suggesting that more than one vesicle was released per synapse during larger barrages. Each data point shows the mean amplitude (z-scored across all events of each synapse) per barrage for geometrically binned barrage sizes.

### Synapses are spatially organized along developing dendrites

In the mature mouse visual cortex, synaptic inputs are functionally clustered along layer 2/3 dendrites, such that neighboring synapses frequently have similar receptive fields (Iacaruso et al., 2017). During development, synaptic inputs are already clustered before eye-opening (Winnubst et al., 2015). Therefore, we explored here the developmental trajectory of dendritic input organization in general and synaptic clustering in particular. First, we noticed – mostly in young neurons – that some dendritic segments contained several synapses, whereas other segments were essentially devoid of active synapses (Fig 5A). To determine whether synapse distribution along the imaged dendrites was indeed patterned rather than random, we compared the distances between each synapse and its nearest distal and proximal neighbors with distances that were estimated after randomly shuffling synaptic locations along the dendrite. We found that the observed inter-synapse distance distribution differed significantly from randomized distributions in younger dendrites (P8-P10; Fig. 5B). In contrast, in older dendrites (P12-P13) synapse distribution was indistinguishable from randomized distributions. We concluded that in younger dendrites synapses accumulated in specific dendritic segments, and that with development the entire dendritic surface became covered by synapses more evenly.

**Fig. 5:**
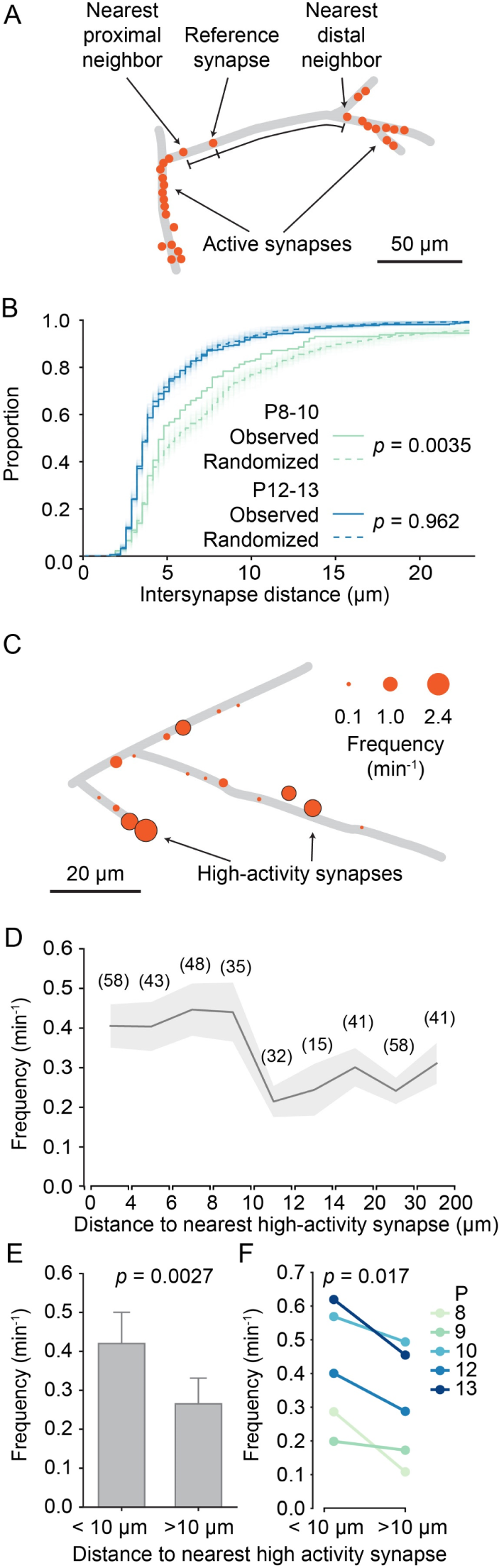
Structured organization of synapses along developing dendrites. A: Graphical representation of layer 2/3 pyramidal cell dendrites at P8 (D2). Each dot shows an active synapse. Several dendritic segments carried a high density of synapses; others did not receive functional synaptic inputs. B: Cumulative distribution of inter-synapse distances between each synapse and their nearest proximal and distal neighbors. The inter-synapse distance distribution of synapses in younger dendrites (P8-10, green) differed significantly from randomized distributions (100 runs are shown as faint lines and their average as dashed line). In older dendrites (P12-13, blue), inter-synapse distributions did not differ from randomized distributions (Kolmogorov-Smirnov tests). C: Graphical representation of layer 2/3 pyramidal cell dendrites at P10 (D5). The diameter of each disc represents the transmission frequency of a synapse. Synapses that were among the 20% most active synapse within their age group (high-activity synapses) are outlined in black. Frequently, highly active synapses were located in close proximity to other highly active synapses. D: The mean transmission frequency of synapses was higher at synapses that were located nearby high-activity synapses. A clear drop in transmission frequency was observed at distances larger than 10 micrometers from the nearest high-activity synapse. In parentheses, the number of synapses averaged for each distance bin are shown. Grey area represents SEMs. E: The mean transmission frequency was higher at synapses that were located within 10 micrometers of a high-activity synapse than at synapses farther away from high-activity synapses (Student’s t-test for independent samples, 2-sided, n = 181 (< 10 µm) and 173 (> 10 µm) synapses. Error bars: SEM). F: Within each age group, synapses that were located within 10 micrometers from an high-activity synapse were more active than those that were located farther away from high-activity synapses (Student’s t-test for paired samples, 2-sided).

Our finding that the distribution of transmission frequencies of individual synapses showed an extended tail (Fig 3F), made us wonder whether those synapses with high transmission frequencies showed any particular spatial distribution as well. We defined high-activity synapses as those in the top 20 activity percentile for each age. In line with our previous observation that spine synapses showed higher transmission rates than shaft synapses (Fig 3G) we found that the majority (76%) of high-activity synapses were located on spines. Furthermore, we observed that highly active synapses occurred frequently near each other (Fig 5C). Therefore, we tested whether there was a relationship between the activity level of a given synapse and the distance to its nearest high-activity neighbor. We discovered that synapses that were located within 10 micrometers from a high-activity synapse were more active than synapses that were more distant to a high-activity synapse (Fig 5D). In fact, neighbors of high-activity synapses were significantly more active than those with larger distances from high-activity synapses (Fig 5E) and this relationship was highly consistent across all ages (Fig 5F).

### Synaptic inputs are organized in functional domains

Having established that high-activity synapses were often surrounded by other high-activity synapses, we defined dendritic segments that contain high-activity synapses and their neighbors as dendritic domains (Fig 6A). Synapses that showed normal activity levels (i.e. non-high-activity synapses) were assigned to domains when they were 10 micrometer or less away from a high-activity synapse. High-activity synapses that were within 10 micrometers of each other were assigned to the same domain, whereas those that were farther away from each other were assigned to separate domains.

**Fig: 6:**
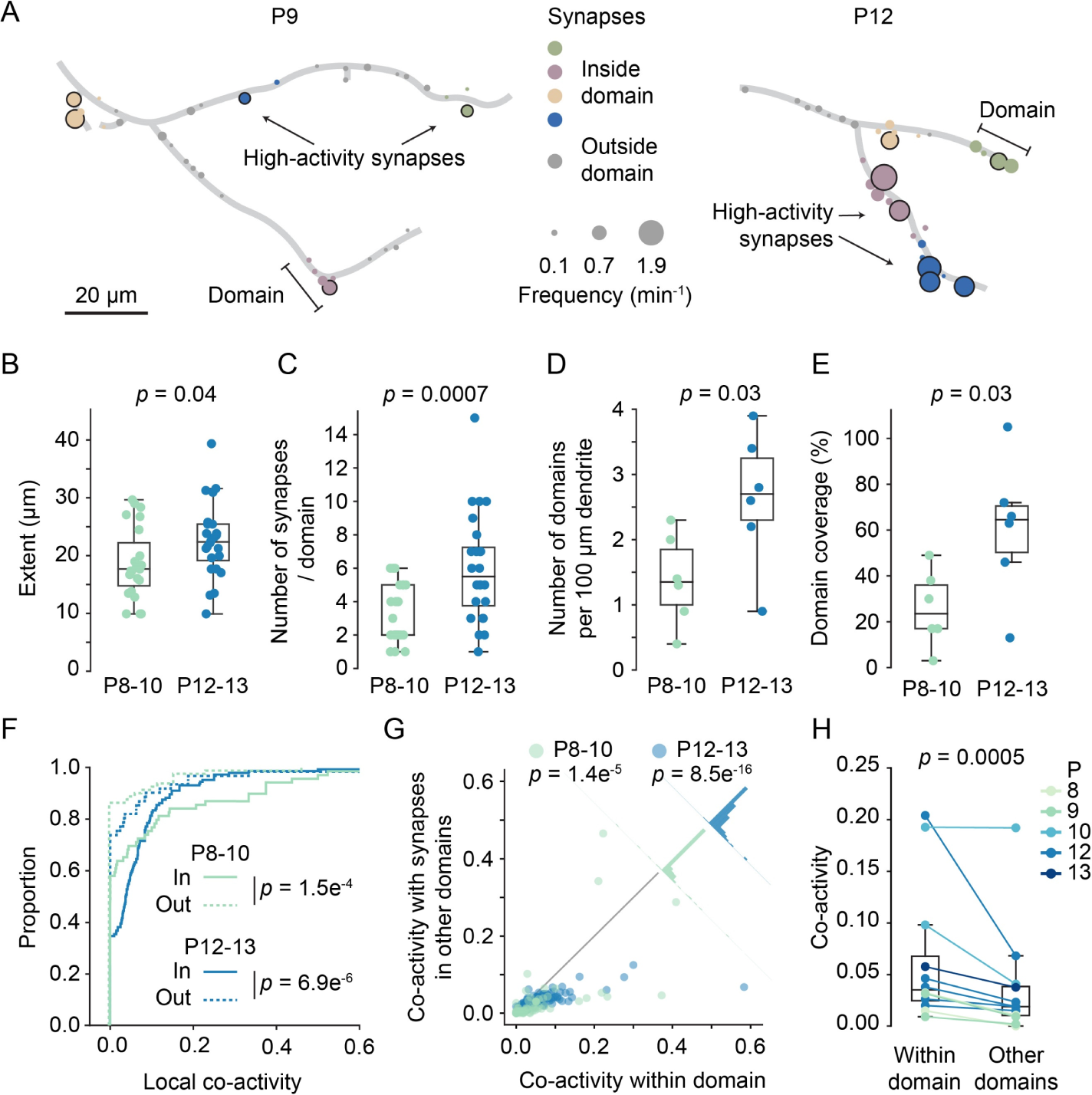
Domains of clustered synapses emerged during the second postnatal week. A: Graphical representation of layer 2/3 pyramidal cell dendrites at P9 (D3) and P12 (D7). Discs represent individual synapses. The disc size indicates synaptic transmission frequency, and high-activity synapses are labeled by black outlines. Individual domains are shown in different colors. Synapses located outside domains are shown in grey. B, C: The extent of domains and the number of synapses increased during development. Each dot represents one domain (Student’s t-test for independent samples, 2-sided, boxes indicate quartiles, and whiskers outline the entire distribution, except outliers). D, E: the density of domains along dendrites and the fraction of dendrite covered by domains (E) increased with age. Each dot represents one dendrite (Student’s t-test for independent samples, 2-sided, boxes indicate quartiles, and whiskers outline the entire distribution, except outliers). F: Cumulative distributions of local co-activity values of all recorded synapses. The local co-activity of synapses inside domains was higher than that of outside-domain synapses (Man-Whitney-U test, 2-sided, n = 213 (In domain) and 173 (Outside domain) synapses. Error bars: SEM) G: Synapses in domains were more co-active with their domain neighbors than with synapses in other domains. Each dot represents the co-activity of a synapse with its neighbors within the domain and its co-activity with synapses in other domains (Wilcoxon signed-rank test). The histograms perpendicular to the equality line show the distribution of differences. H: For each experiment, the co-activity of synapses within domains was higher than their co-activity with synapses in other domains (Wilcoxon signed-rank test).

Assessing domain features across development revealed that their extent increased slightly (P8-10: 18.7 ± 6.1 µm; P12-13: 22.5 ± 6.4 µm). Moreover, both the number of synapses per domain (P8-10: 3.2 ± 1.7; P12-13: 6.0 ± 3.2) as well as the density of domains along dendrites (P8-10: 1.4 ± 0.7; P12-13: 2.6 ± 1.0 per 100 µm dendrite) approximately doubled during the second postnatal week (Fig 6B-D). The fraction of dendritic length covered by domains had increased significantly by the end of the second postnatal week (Fig 6E; P8-10: 27.2 ±16.2%; P12-13: 61.3 ± 29.4%, mean ± STD). Finally, the percentage of synapses within domains increased with age as well (P8-10: 46.2 ± 7.4%, n = 6 dendrites; P12-13: 72 ± 23.2%, n = 6 dendrites, mean ± STD; p = 0.026, t-test).

To relate dendritic domains (i.e. segments surrounding high-activity synapses) to synapse clusters (i.e. neighboring synapses with similar activity patterns), we compared local co-activity between synapses both inside and outside of dendritic domains. For each synapse, we determined a co-activity value by dividing the number of times this synapse was co-active with any synapse in the comparison group, e.g. its neighbors, by the number of times this synapse was active in total, normalized by the number of synapses in the comparison group. Next, we compared the local co-activity of synapses within domains to the local co-activity of synapses outside domains, and found that domain synapses showed much higher local co-activity in both younger (in: 0.08 ± 0.02; out: 0.02 ± 0.01; mean ± SEM) and older animals (Fig 6F; in: 0.06 ± 0.01; out: 0.04 ± 0.01; mean ± SEM). Thus, synapses within a domain were more synchronized with their neighbors than synapses that were not part of a domain.

This finding raised the question whether dendritic domains are also functionally distinct from each other such that patterns of synaptic inputs received by one domain differed from the patterns received by the other domains on the same dendrite. To address this possibility, we quantified the co-activity of individual synapses with synapses from within their domain and compared that with their co-activity with synapses from other domains. We found that co-activity within domains was consistently higher than co-activity between synapses from different domains across all imaged dendrites (Fig 6G-H). These results demonstrate that synapses with similar input patterns emerge in distinct dendritic domains. Furthermore, they suggest that clustered synaptic inputs in the mouse visual cortex (Iacaruso et al., 2017; Wilson et al., 2016; Winnubst et al., 2015) are confined to developmentally emerging distinct dendritic domains, and do not represent a smooth gradient of input patterns along the dendrite.

### Local synaptic synchronicity correlates with synaptic activity changes

We then asked how uniform synaptic inputs became sorted into domains. Since we had discovered previously that local co-activity drives plasticity to cluster neighboring synapses (Niculescu et al., 2018; Winnubst et al., 2015) and found here, that synapses within domains were more synchronized with their neighbors than synapses located outside domains (Fig. 6F), we examined whether co-activity of neighboring synapses within domains predicted changes in their activity levels. To quantify changes in activity over time, we determined the Mann-Kendall score (Hussain and Mahmud, 2019) for each synapse. This score is positive for monotonic increases, negative for decreases, and zero in the absence of changes. Thus, synapses that increased in transmission frequency across recordings of an experiment yielded positive scores, those that underwent synaptic depression (i.e. decreased in transmission frequency) yielded negative scores, and those that maintained activity levels scored at zero (Fig 7A). We found that the Mann-Kendall score was more negative outside than inside domains, indicating that on average synapses outside domains tended to decrease in activity over time whereas those inside domains showed a more positive activity trajectory (Fig 7B). Next, we compared each synapse’s Mann-Kendall score of synaptic plasticity with its local co-activity within 10 micrometers proximally and distally along the dendrite and revealed a significant positive correlation (Fig 7A, C), demonstrating that synapses that were synchronized with their neighbors became stabilized or more active compared to those that were out of sync with their neighbors.

**Fig. 7:**
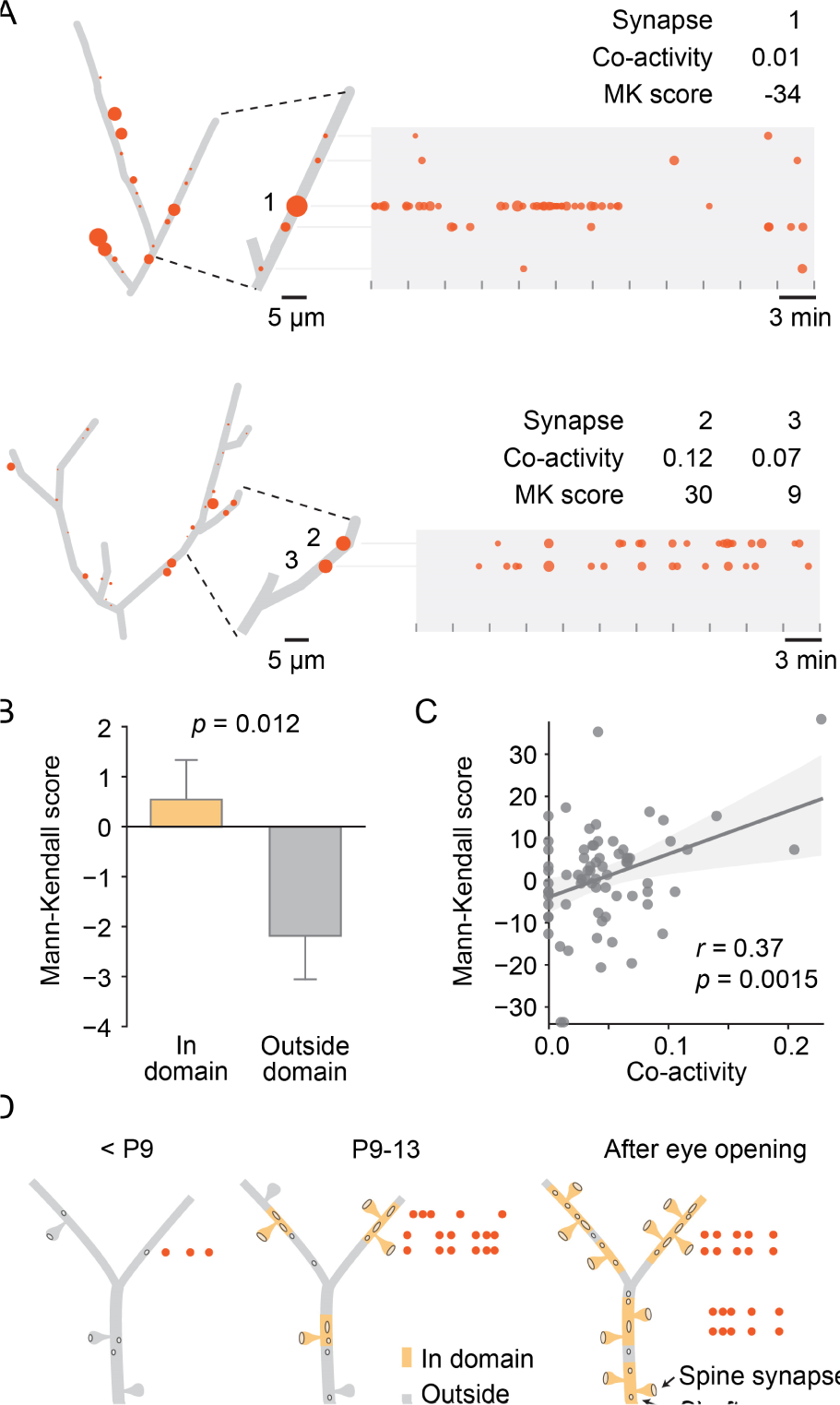
Local co-activity of synapses inside dendritic domains correlated with synaptic potentiation. A: Graphical representation of two layer 2/3 pyramidal cell dendrites at P12 (D5, D9). Left: Red discs on each dendrite represent active synapses, and their size shows relative activity levels. Activity of synapse (1) was mostly out-of-sync with its neighbors as indicated by a low co-activity score. This synapse was very active at the beginning of the experiment and stopped transmitting towards the end, resulting in a low Mann-Kendall (MK) score, which indicates a monotonic decrease in synaptic activity. Synapses (2) and (3) showed higher local co-activity and increased in activity over time as evidenced by a positive MK score. Tick marks indicate durations of individual 3 minute recordings. Red dots in the time plots represent individual synaptic transmission events. The size of each data point indicates the relative amplitude of each event. B: The Mann-Kendall score, a measure of monotonic changes in synaptic transmission frequency, was higher for domain synapses than synapses located outside domains. (Student’s t-test for independent samples, 2-sided, n = 213 (In domain) and 173 (Outside domain) synapses from all ages. Error bars: SEM) C: The change in transmission frequency of a synapse was correlated with its local co-activity. Each dot represents one synapse with a transmission frequency of at least 0.6 min^-1^ (Linear regression). D: Model for the establishment of domains of clustered synaptic inputs in layer 2/3 neurons of the developing visual cortex. Until P8, only a few active synapses are scattered across the developing dendrites. During the second postnatal week, synapses become sorted into distinct domains where in-sync synapses become stabilized or potentiated, and desynchronized synapses undergo synaptic depression. By eye-opening, most dendrites are covered by domains of synapses that are more co-active with each other than with synapses that are located in other domains.

Together, these findings showed that visual cortex neurons established functional synapses at a very high rate during the week leading up to eye opening and suggest that developing synapses became organized into functional, spatially distinct domains based on the synchronicity with their neighbors (Fig 7D).

## Discussion

The fundamental idea that local processing and plasticity in dendrites is essential for high-level computations of the brain (Mel et al., 2017; Poirazi et al., 2003; Poirazi and Mel, 2001) has gained tremendous traction. Many recent studies show that synaptic inputs to excitatory neurons in the neocortex and hippocampus are clustered, such that synapses representing similar stimulus features are located near each other. However, when and how structured synaptic inputs assemble during development has been unclear. Here, we show that synaptic inputs to layer 2/3 pyramidal cells in the mouse visual cortex are structurally and functionally organized in distinct dendritic domains already during the second postnatal week, i.e. before eye opening and detailed visual input. Furthermore, we find that local co-activity within dendritic domains correlates with synaptic activity changes, indicating that synapses are sorted into distinct functional dendritic domains through plasticity mechanisms driven by spontaneous network activity (Fig 7D).

We obtained insight into the spatial organization of synaptic inputs onto layer 2/3 pyramidal cell dendrites by mapping synaptic transmission during spontaneously occurring network activity *in vivo*. Hence, the observed synaptic activity is the result of an interaction between the firing patterns of presynaptic neurons and the features of the individual synapses on the monitored neuron. Therefore, it is important to understand which properties of synaptic inputs we can infer from the recorded input patterns. Firstly, we observed that the distribution of synaptic transmission frequencies is very long-tailed: most synapses show only rare transmission events, whereas roughly 20 percent of synapses are highly active. In contrast, the largest population of input neurons (i.e. other layer 2/3 pyramidal cells) show very homogeneous firing activity during neonatal spontaneous network events where all neurons participate at a similar rate (Rochefort et al., 2009; Siegel et al., 2012). Consequently, the distribution we observed here for individual synapses is most likely determined by release probability differences between individual presynaptic terminals along the dendrite. Furthermore, our current findings are congruent with our previous results which show that the transmission frequency of individual synapses is highly plastic and regulated over a large range by local synaptic interactions and a push-pull mechanism between mature BDNF and proBDNF in pyramidal neurons (Niculescu et al., 2018; Winnubst et al., 2015). Thus, transmission frequency is a synapse feature of large dynamic range that is most likely harnessed by the developing circuit to adjust connection strength in response to fine-scale activity patterns. This conclusion is supported by a previous study reporting that differences in cortico-cortical synaptic connections are mostly based on differences in synaptic release probabilities, and that these release probabilities change in response to altered input patterns (Markram et al., 1997). Furthermore, the amplitude of calcium transients in axonal boutons varies over an order of magnitude, most likely representing a similar degree of variability in release probability, in juvenile rat layer 2/3 pyramidal cells (Koester and Sakmann, 2000).

How many independent inputs, i.e. from unique presynaptic neurons, become connected to one dendritic domain? We observed a mean increase from three to six synapses per domain during the second postnatal week. In the adult, connected cortical neurons typically connect through fewer than ten synapses, most of which are located on different dendrites (Feldmeyer et al., 2006; Markram et al., 1997; Winfield et al., 1981). However, axons that establish more than one synapse with individual dendritic segments have been observed as well. For example, in the mature somatosensory cortex, approximately ten percent of synapses are redundant, i.e. dendritic segments receive another synapse (or more) from the same axon (Kasthuri et al., 2015). At the ages investigated in the present study, synapse density is lower than in the mature sensory cortex and redundant synapses most likely rarer. Nevertheless, our data set probably contains redundant synapses, too. Interestingly, full connectomic reconstruction of mature cortical dendrites and their presynaptic axons suggested that synaptic partner selection based on coincident activity may be responsible for the formation of multiple synapses of one axon with the same dendritic segment (Kasthuri et al., 2015), indicating that multiple innervation may be generated through the here described plasticity mechanism that helps shaping domains of co-active synapses.

In the mature visual system, V1 pyramidal cell synaptic inputs are clustered, such that similar stimulus features are represented by neighboring synapses (Iacaruso et al., 2017; Wilson et al., 2016). This input organization is thought to underlie local dendritic computations that maximize the computational power of cortical pyramidal neurons (Poirazi and Mel, 2001). In fact, perturbing local dendritic integration has been shown to decrease the sensitivity or selectivity of cortical neurons to stimuli of different modalities (Lavzin et al., 2012; Palmer et al., 2014; Smith et al., 2013; Xu et al., 2012). Which stimulus features may be encoded by synaptic domains established before eye opening? In the adult, neighboring synapses are more likely to share receptive field properties than more distant synapses. For example, in ferret V1 layer 2/3 neurons, synapses are clustered according to orientation preference (Wilson et al., 2016), whereas in mouse V1 layer 2/3 neurons, synapses are clustered based on receptive field localization within the visual field, as well as receptive field similarity, but not on orientation preference (Iacaruso et al., 2017). Thus, different sensory input features can be arranged in neighborhoods along dendrites, but exactly which features are clustered differs between species. In the retina, spontaneously generated activity travels in waves with specific directions as well as wavefronts of defined orientations. In addition, neighboring neurons are frequently co-active during wave propagating (Feller et al., 1997; Meister et al., 1991). Hence, retinal waves encode retinotopy as well as direction and orientation selectivity. Since wave activity in retinal ganglion cells is transmitted along the central visual pathways from the retina to the visual cortex (Ackman et al., 2012; Hanganu et al., 2006; Siegel et al., 2012), retinal waves can shape dendritic domains based on neighborhood relationships of retinal ganglion cells representing location in visual space, and orientation and direction selectivity (Kirchner and Gjorgjieva, 2021). Finally, our observation that functional synapses form in distinct domains along dendrites, whereas other dendritic segments are devoid of synapses at the beginning of the second postnatal week, indicates that synapses cluster in defined domains rather than distributing evenly along dendrites.

Together, patterned spontaneous activity can be sufficient to sort synaptic inputs into domains to establish discrete computational modules along dendrites to prepare the visual cortex for processing with high sensitivity and selectivity before the eyes open. This “first guess” will be refined further by visual inputs after eye opening, based on the statistics of sensory inputs in the actual environment.

## Methods

### Animals

All experimental procedures were approved by the Institutional Animal Care and Use Committee of the Royal Netherlands Academy of Arts and Sciences. We used 11 C57BL/6J mouse pups between P8 and P13.

### Plasmids

For *in utero* electroporation, GCaMP6s (Addgene plasmid 40753; Douglas Kim; Chen et al., 2013) was cloned into pCAGGS and used in combination with DsRed Express in pCAGGS (Winnubst et al., 2015, gift from Christiaan Levelt). Cells co-expressed the fluorescent protein DsRed for somatic targeting and structural information.

### In Utero Electroporation

Constructs were introduced through in utero electroporation at E16.5. Pyramidal neurons in layer 2/3 of the visual cortex were transfected with GCaMP6s (2 mg/ml) and DsRed (0.5-2 mg/ml) at E16.5 using in utero electroporation (Winnubst et al., 2015). Pregnant mice were anesthetized with isoflurane and a small incision (1.5–2 cm) was made in the abdominal wall. The uterine horns were removed from the abdomen, and DNA (1 µl) was injected into the lateral ventricle of embryos using a sharp glass pipette. Voltage pulses (five square wave pulses, 30 V, 50 ms duration, 950 ms interval, custom-built electroporator) were delivered across the brain with tweezer electrodes covered in conductive gel. Uterine horns were rinsed with warm saline solution and carefully returned to the abdomen, after which the muscle and skin were sutured.

### *In vivo* electrophysiology and calcium imaging

The surgery and stabilization for the *in vivo* calcium imaging experiments were performed as described previously (Siegel et al., 2012; Winnubst et al., 2015). Animals were anesthetized with 2% isoflurane, which was reduced to 0.7%–1% after surgery. We previously reported that although this low level of anesthesia does reduce the frequency of spontaneous network events, relative to that seen in awake animals, it does not change the basic properties of spontaneous network activity, such as participation rates and event amplitudes (Siegel et al., 2012). Furthermore, we found previously that synaptic inputs are organized very similarly *in vivo* under light anesthesia and in slice cultures (Winnubst et al., 2015). Together, these observations indicate that the activity patterns investigated here are not or only slightly affected by low-level anesthesia. Glass electrodes (4.5–6 MΩ) were fluorescently coated with BSA-Alexa 594 to allow targeted whole-cell recordings (Sasaki et al., 2012). To visualize synaptic inputs, action potentials were blocked with QX314 in the intracellular solution (120 mM CsMeSO_3_, 8 mM NaCl, 15 mM CsCl_2_, 10 mM TEA-Cl, 10 mM HEPES, 5 mM QX-314 bromide, 4 mM MgATP, and 0.3 mM Na-GTP; Takahashi et al., 2012; Winnubst et al., 2015). Currents were recorded in voltage clamp mode at 10 kHz and filtered at 3 kHz (Multiclamp 700b; Molecular Devices). To facilitate the detection of synaptic calcium transients, neurons were depolarized to –30 mV to increase NMDA receptor activation (Takahashi et al., 2012; Winnubst et al., 2015). When switching to –30 mV, the baseline fluorescence of the patched neurons increased, but not of the neighboring neurons, allowing unequivocal identification of all dendrites in the imaged volume that belonged to the patched neuron (Fig S3). No correction was made for the liquid junction potential. Barrages were identified as fast rising inward currents of at least 10 pA, followed by a return to baseline, which consisted of multiple synaptic currents and were separated by at least three seconds.

### Image acquisition

*In vivo* calcium imaging was performed on a Nikon (A1R-MP) with a 0.8/16x water-immersion objective and a Ti:Sapphire laser (Chameleon II, Coherent). Dendrites were imaged at 5-15 Hz with a pixel size of 0.32 µm in single planes or small stacks with planes spaced 1.5 – 2 µm. We recorded the movement signal of the scan mirrors to synchronize calcium imaging and electrophysiology.

### Image processing

To remove drift and movement artifacts from each recording, we performed drift correction using NoRMCorre (Pnevmatikakis and Giovannucci, 2017). Each recording was aligned to the first recording in the series to remove any movements between recording sessions. Delta-F stacks were made using the average fluorescence per pixel as baseline. Local synaptic calcium transients were clearly detectable in the delta-F stacks.

Regions of interest (ROIs), representing putative synapses, were hand-drawn using ImageJ (NIH) at sites where spines were visible and at clearly observed activity sites (Fig. S4). Finally, all remaining dendritic segments were filled with ROIs to ensure that no sites were missed. Putative synaptic transmission events were automatically identified across all ROIs as mean fluorescence increases that were at least two-fold higher than the noise level. These putative signals were then manually reviewed using the full image recordings. Signals were rejected, if they were shorter than two frames or coincided with movement. Furthermore, we ensured that there was a clearly visible local maximum at the location of the synapse and that it was stable for the duration of the signal. Automated transient detection and further data processing was performed using custom-made Matlab (MathWorks) and Python software (Python Software Foundation).

### Identification of spine and shaft synapses

We identified spine and shaft synapses as follows. Synaptic calcium transients that occurred in small appendages from the dendrite were defined as transmission events at spine synapses (e.g. Fig 2A and Fig S5A). Synaptic calcium transients located in the center of the dendrite, but at sites that showed small, sharply confined areas of increased fluorescence and where 3D reconstructions suggested that a spine may be located above or below the dendrite in the z-dimension were defined as spine transmission events as well (e.g. S5B). Synaptic calcium transients that occurred in the center of “smooth” dendritic stretches (without clearly defined spots of increased brightness) were identified as transmission events at shaft synapses (e.g. Fig 2B and Fig S5A). Since the z-resolution of 2-photon microscopy is lower in the z-dimension than in xy, we may have misidentified spine synapses as shaft synapses in a few cases.

### Statistical Analysis

One dendrite was imaged from each animal except once, when two dendrites were imaged in a single cell in a P12 animal. We used only synaptic sites that were active with a transmission frequency of at least 0.03 per minute. We tested whether the localization of functional synapses was structured or random along the dendrite. To establish randomized distributions of synapse positions, we first assigned “empty” positions between the positions of actual functional synapses such that the entire imaged dendrite was covered with real or empty positions at the typical minimal inter-synapse distance (4 ± 1 µm). Next we determined all pair-wise distances between all synapses and all empty locations along dendrites using the Shortest Path with Obstacle Avoidance function of Matlab. Then we shuffled all functional synapses across the real and empty positions randomly. For each shuffle and the observed localizations, we determined the distances of each synapse to its nearest proximal and distal neighbors. Finally we compared the results of 100 shuffles with the observed distribution using a Kolmogorov Smirnov test.

The co-activity of a given synapse (e.g. with its neighbors within 10 micrometers, with its neighbors in the same domain or with synapses in other domains) was determined by dividing the number of observed transmission events at the observed synapse, by the number of transmission events of the comparison group (e.g. its neighbors) during the same barrages when the observed synapse was active. This number was divided by the number of synapses in the comparison group to normalize to the size of the comparison group. We defined co-activity based on co-transmission during the same barrage, because we had previously discovered that co-activity during barrages is crucial for local synaptic plasticity (Winnubst et al., 2015).

Domains were defined as dendritic segments that contained at least one high-activity synapse (see Results section for definition). If the distance between two high-activity synapses was ten micrometers or less, both were assigned to the same domain. Normal synapses (those that were not high-activity synapses) were assigned to a domain, if their distance to the nearest high-activity synapse was ten micrometers or less. Normal synapses located between two high-activity synapses from distinct domains with 20 micrometer or less distance were assigned to the domain of the closer high-activity synapses. The extent of dendritic domains was determined as the distance between its most distant members. The length was padded at the edges with five micrometers from a high-activity synapse, if there was no normal member within this distance. Thus single high-activity synapse domains had a length of ten micrometers.

For normally distributed data and data with relatively low sample numbers (6-8), where normality tests are underpowered, we used paired and unpaired, 2-tailed Student’s t-tests as recommended (de Winter, 2013). For data that were not normally distributed, we used non-parametric statistical comparisons, Man-Whitney-U tests for independent samples and Wilcoxon signed-rank tests for paired data. To test whether investigated parameters changed with age, we performed linear regression analyses. To test the differences between distributions, we used a Kolmogorov-Smirnov test.

## Author contributions

Conceptualization: A.H.L., J.E.C. and C.L.; Investigation: A.H.L., J.E.C.; Analysis: A.H.L., J.E.C. and C.L.; Writing: A.H.L., J.E.C. and C.L..

## Acknowledgments

We thank Jan Kirchner and Julijana Gjorgjieva for discussions, Christiaan Levelt for sharing plasmids, and Helmut Kessels, Alexander Heimel, Wei Wei, David Cabrera, Tamara Buijs, and Julijana Gjorgjieva for critically reading this manuscript. This work was supported by grants of the Netherlands Organization for Scientific Research (NWO, ALW Open Program grants, no. 819.02.017, 822.02.006 and ALWOP.216; ENW Open Competition grant no. OCENW.KLEIN.535, ALW Vici, no. 865.12.001), ZonMW (Top grant no. 9126021) and the “Stichting Vrienden van het Herseninstituut” (all CL.).

## Supplementary Figures

**Fig. S1.**
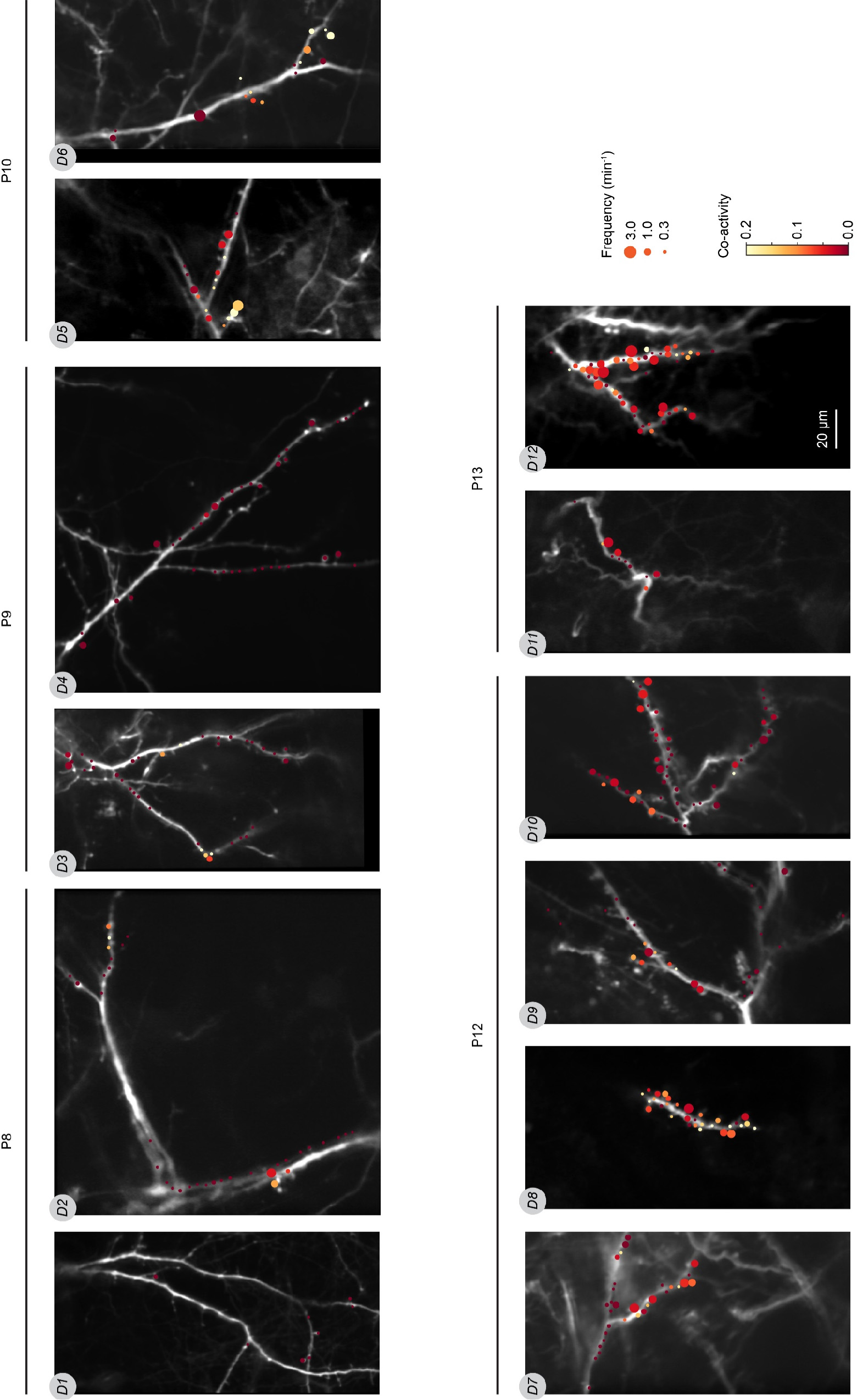
Overview of all imaged dendrites and synapses. Each synapse is represented as a disc whose size and color indicate transmission frequency and local co-activity, respectively. D1-D12 are each dendrite’s identifier. The scale bar applies to all images.

**Fig. S2.**
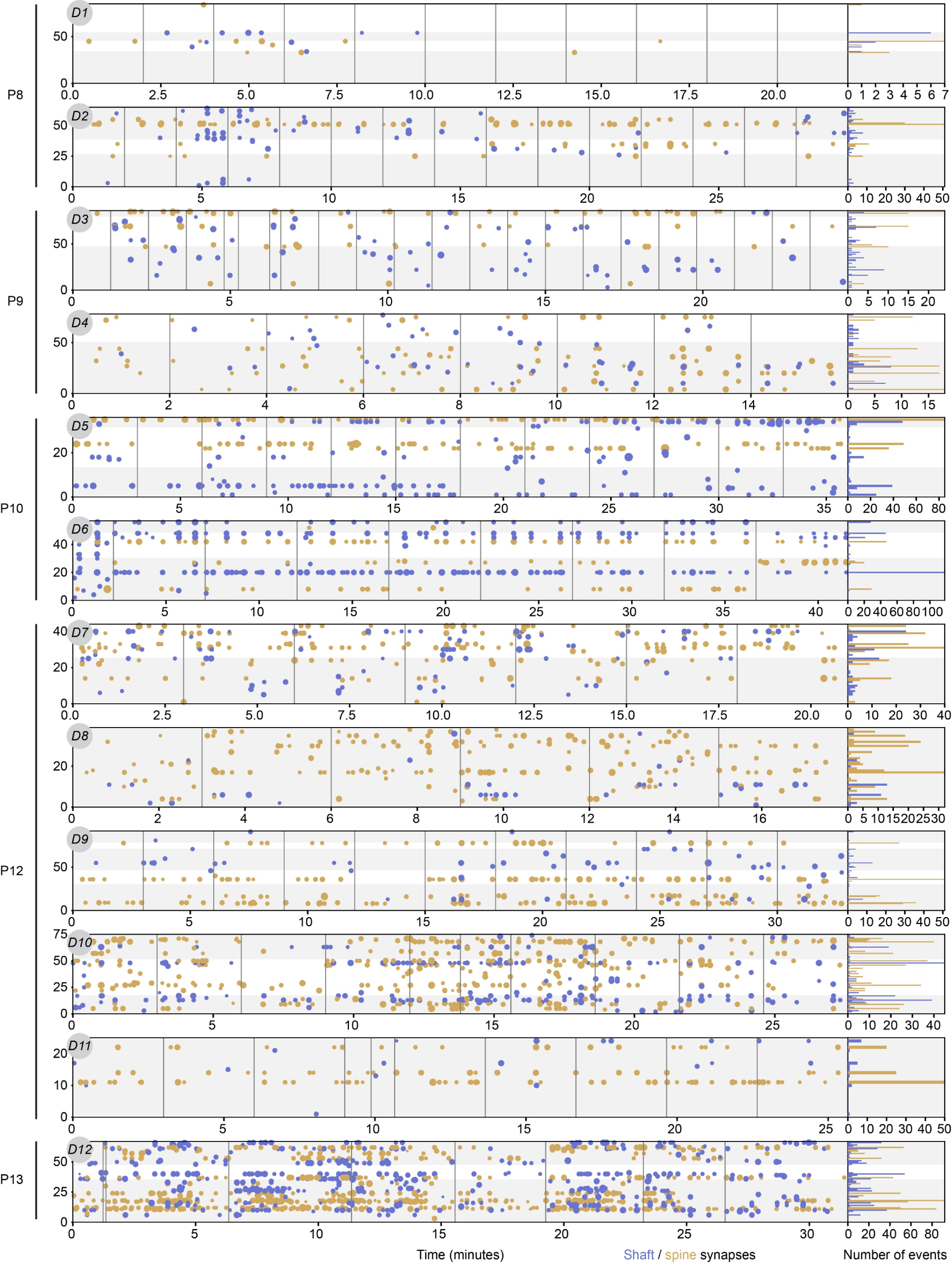
Overview of all recorded synaptic transmission events across all imaged dendrites. Y-axes show synaptic position along dendrites. White and grey shaded areas represent individual dendritic segments. Each synaptic transmission event is shown as a disc in yellow (spines) or blue (shaft synapses). Disc size indicates the relative amplitude of each event (z-scored for each synapse). Bar plots on the right show the number of transmission events detected at each synapse. Individual dendrites are shown in the same sequence as in Fig. S1.

**Fig. S3.**
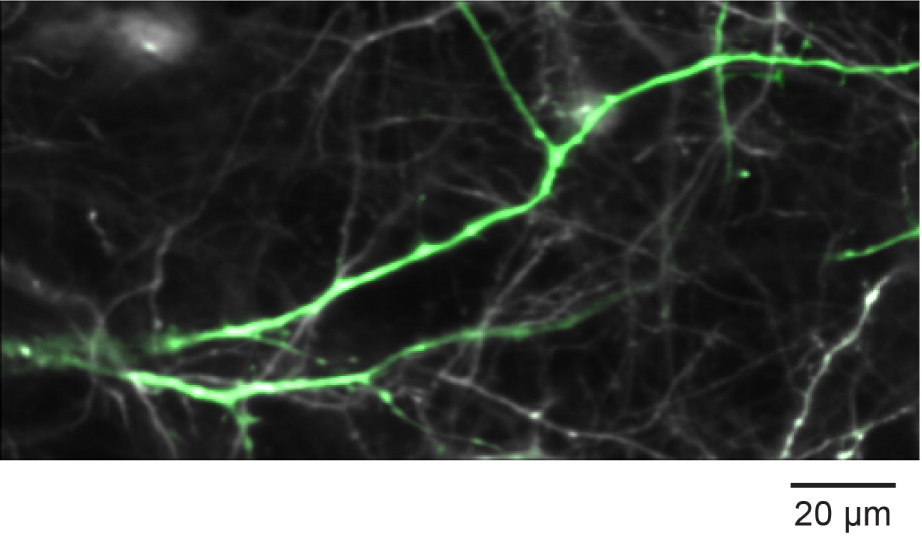
Identification of the patched neuron and its dendrites. Composite of the DSred channel (grey) and the GCaMP6s channel (green) after switching the holding potential to – 30 mV of a layer 2/3 pyramidal neuron at P8 (D1). The depolarization triggers a transient calcium increase in the patched neuron, which labels its dendrites (green) clearly above the background (grey processes).

**Fig. S4.**
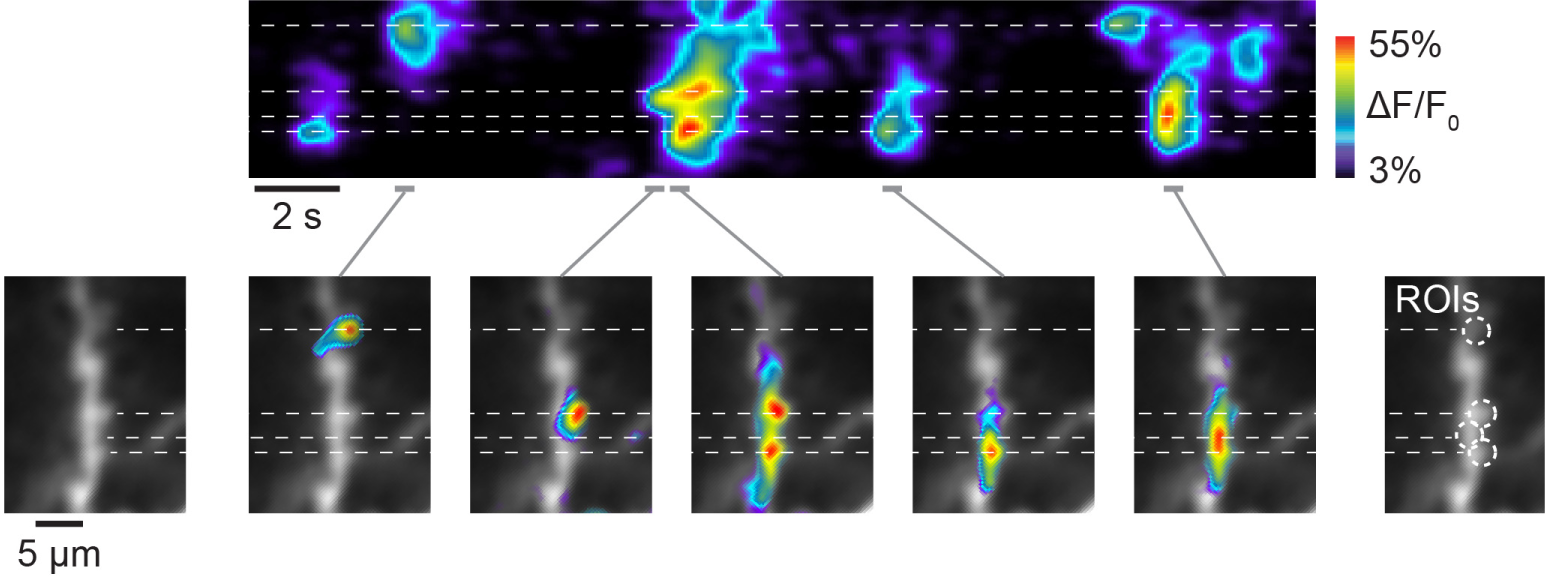
ROI selection. Synaptic calcium transients in a dendrite of a layer 2/3 pyramidal neuron at P13 (D12). Top: occurrence of synaptic calcium transients in the dendritic stretch shown on the left. Stippled lines indicate the position of several transients. Bottom: Spatial representation of the events shown above at the time points indicated by the grey bars. Circles on the right indicate ROIs selected based on the calcium transients shown here.

**Fig. S5.**
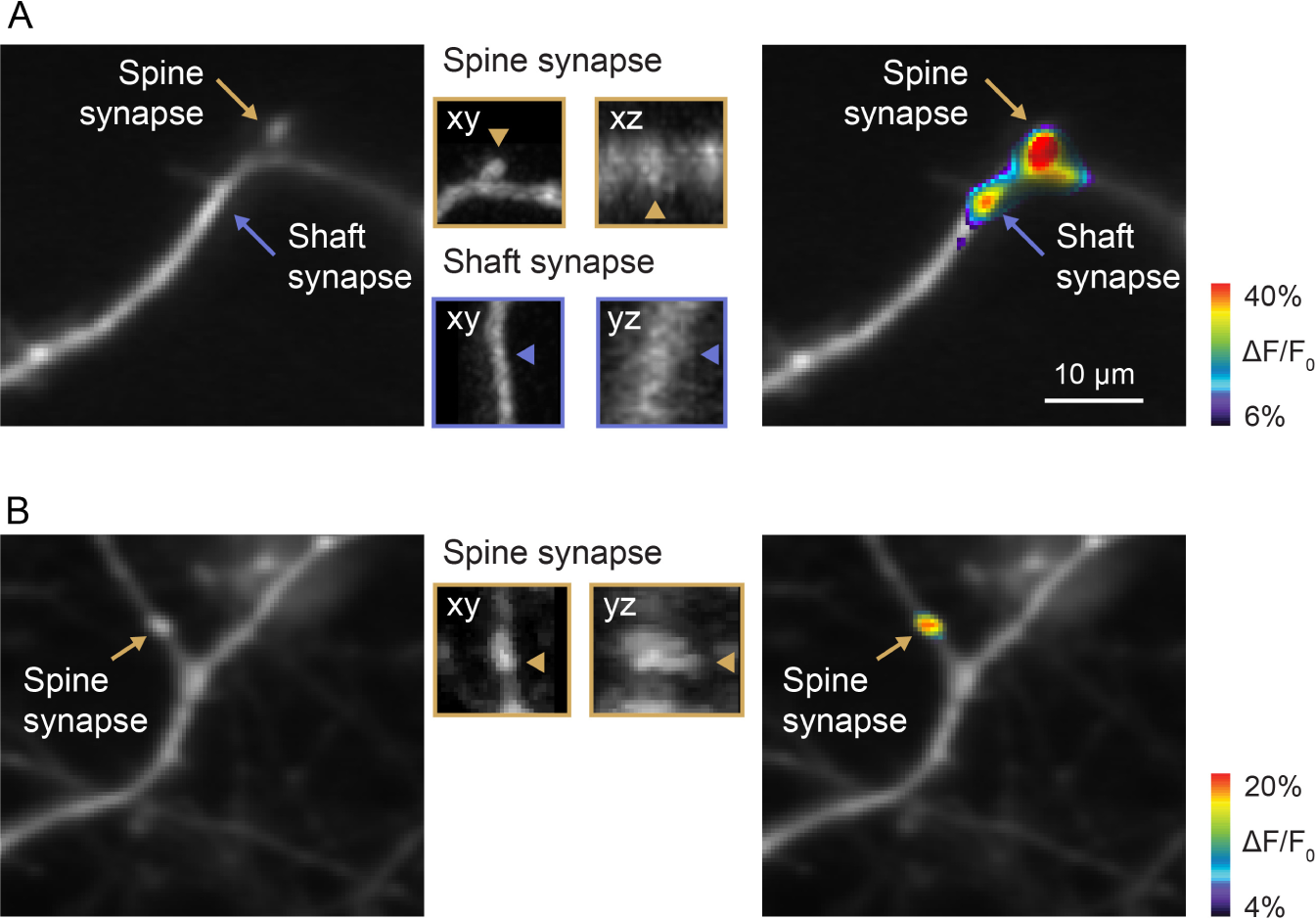
Identification of shaft and spine synapses. A: Dendrite of a layer 2/3 pyramidal neuron at P9 (D3). Left: Arrows indicate the positions of a spine (yellow) and a shaft synapse (blue). Middle: xy– and xz/yz-views show the dendritic segments around these synapses acquired in a z-stack before the here shown time-lapse recording. Arrow heads point to the same synapses as in the left panel. Right: Synaptic transmission events at these synapses. B: Dendrite of a layer 2/3 pyramidal neuron at P8 (D1). The arrow points to a putative spine synapse located in the axis of the dendritic shaft. xy– and yz-views show the dendritic segment around this synapse acquired in a z-stack before the here shown time-lapse recording. The arrowheads points to a putative spine below the dendrite in the image plane. Right: synaptic transmission event at this synapse.

## References

1. Ackman JB, Burbridge TJ, Crair MC. 2012. Retinal waves coordinate patterned activity throughout the developing visual system. Nature 490:219–225.

2. Berry KP, Nedivi E. 2017. Spine dynamics: Are they all the same? Neuron 96:43–55. doi:10.1016/j.neuron.2017.08.008

3. Blue ME, Parnavelas JG. 1983. The formation and maturation of synapses in the visual cortex of the rat. I. Qualitative analysis. J Neurocytol 12:599–616.

4. Branco T, Hausser M. 2011. Synaptic integration gradients in single cortical pyramidal cell dendrites. Neuron 69:885–892.

5. Chen T-W, Wardill TJ, Sun Y, Pulver SR, Renninger SL, Baohan A, Schreiter ER, Kerr RA, Orger MB, Jayaraman V, Looger LL, Svoboda K, Kim DS. 2013. Ultrasensitive fluorescent proteins for imaging neuronal activity. Nature 499:295–300.

6. Chen X, Leischner U, Rochefort NL, Nelken I, Konnerth A. 2011. Functional mapping of single spines in cortical neurons in vivo. Nature 475:501–505.

7. Cichon J, Gan W-B. 2015. Branch-specific dendritic Ca2+ spikes cause persistent synaptic plasticity. Nature 520:180–185. doi:10.1038/nature14251

8. Cline H. 2003. Sperry and Hebb: oil and vinegar? Trends Neurosci 26:655–661.

9. De Felipe J., Marco P, Fairen A, Jones EG. 1997. Inhibitory synaptogenesis in mouse somatosensory cortex. Cereb Cortex 7:619–634.

10. de Winter J. 2013. Using the Student’s t-test with extremely small sample sizes. Pract Assess Res Eval 18.

11. Feldmeyer D, Lübke J, Sakmann B. 2006. Efficacy and connectivity of intracolumnar pairs of layer 2/3 pyramidal cells in the barrel cortex of juvenile rats. J Physiol 575:583–602. doi:10.1113/jphysiol.2006.105106

12. Feller MB, Butts DA, Aaron HL, Rokhsar DS, Shatz CJ. 1997. Dynamic processes shape spatiotemporal properties of retinal waves. Neuron 19:293–306.

13. Fu M, Yu X, Lu J, Zuo Y. 2012. Repetitive motor learning induces coordinated formation of clustered dendritic spines in vivo. Nature 483:92–95.

14. Hanganu IL, Ben Ari Y, Khazipov R. 2006. Retinal waves trigger spindle bursts in the neonatal rat visual cortex. J Neurosci 26:6728–6736.

15. Harnett MT, Makara JK, Spruston N, Kath WL, Magee JC. 2012. Synaptic amplification by dendritic spines enhances input cooperativity. Nature 491:599–602.

16. Hedrick NG, Lu Z, Bushong E, Singhi S, Nguyen P, Magaña Y, Jilani S, Lim BK, Ellisman M, Komiyama T. 2022. Learning binds new inputs into functional synaptic clusters via spinogenesis. Nat Neurosci 25:726–737. doi:10.1038/s41593-022-01086-6

17. Helmchen F, Svoboda K, Denk W, Tank DW. 1999. In vivo dendritic calcium dynamics in deep-layer cortical pyramidal neurons. NatNeurosci 2:989–996.

18. Hussain MM, Mahmud I. 2019. pyMannKendall: a python package for non parametric Mann Kendall family of trend tests. J Open Source Softw 4:1556. doi:10.21105/joss.01556

19. Iacaruso MF, Gasler IT, Hofer SB. 2017. Synaptic organization of visual space in primary visual cortex. Nature 547:449–452. doi:10.1038/nature23019

20. Jia H, Rochefort NL, Chen X, Konnerth A. 2010. Dendritic organization of sensory input to cortical neurons in vivo. Nature 464:1307–1312.

21. Ju N, Li Y, Liu F, Jiang H, Macknik SL, Martinez-Conde S, Tang S. 2020. Spatiotemporal functional organization of excitatory synaptic inputs onto macaque V1 neurons. Nat Commun 11:697. doi:10.1038/s41467-020-14501-y

22. Kasai H, Ziv NE, Okazaki H, Yagishita S, Toyoizumi T. 2021. Spine dynamics in the brain, mental disorders and artificial neural networks. Nat Rev Neurosci 1–16. doi:10.1038/s41583-021-00467-3

23. Kasthuri N, Hayworth KJ, Berger DR, Schalek RL, Conchello JA, Knowles-Barley S, Lee D, Vázquez-Reina A, Kaynig V, Jones TR, Roberts M, Morgan JL, Tapia JC, Seung HS, Roncal WG, Vogelstein JT, Burns R, Sussman DL, Priebe CE, Pfister H, Lichtman JW. 2015. Saturated Reconstruction of a Volume of Neocortex. Cell 162:648–661. doi:10.1016/j.cell.2015.06.054

24. Kerlin A, Boaz M, Flickinger D, MacLennan BJ, Dean MB, Davis C, Spruston N, Svoboda K. 2019. Functional clustering of dendritic activity during decision-making. eLife 8:e46966. doi:10.7554/eLife.46966

25. Kirchner JH, Gjorgjieva J. 2021. Emergence of local and global synaptic organization on cortical dendrites. Nat Commun 12:4005. doi:10.1038/s41467-021-23557-3

26. Koester HJ, Sakmann B. 2000. Calcium dynamics associated with action potentials in single nerve terminals of pyramidal cells in layer 2/3 of the young rat neocortex. J Physiol 529:625–646. doi:10.1111/j.1469-7793.2000.00625.x

27. Larkum ME. 2022. Are Dendrites Conceptually Useful? Neuroscience 489:4–14. doi:10.1016/j.neuroscience.2022.03.008

28. Larkum ME, Nevian T. 2008. Synaptic clustering by dendritic signalling mechanisms. Curr Opin Neurobiol 18:321–331.

29. Lavzin M, Rapoport S, Polsky A, Garion L, Schiller J. 2012. Nonlinear dendritic processing determines angular tuning of barrel cortex neurons in vivo. Nature 490:397–401.

30. Lohmann C, Kessels HW. 2014. The developmental stages of synaptic plasticity. J Physiol 592:13–31. doi:10.1113/jphysiol.2012.235119

31. Losonczy A, Magee JC. 2006. Integrative Properties of Radial Oblique Dendrites in Hippocampal CA1 Pyramidal Neurons. Neuron 50:291–307.

32. Major G, Larkum ME, Schiller J. 2013. Active properties of neocortical pyramidal neuron dendrites. Annu Rev Neurosci 36:1–24. doi:10.1146/annurev-neuro-062111-150343

33. Makara JK, Magee JC. 2013. Variable Dendritic Integration in Hippocampal CA3 Pyramidal Neurons. Neuron 80:1438–1450. doi:10.1016/j.neuron.2013.10.033

34. Makino H, Malinow R. 2011. Compartmentalized versus Global Synaptic Plasticity on Dendrites Controlled by Experience. Neuron 72:1001–1011.

35. Markram H, Lubke J, Frotscher M, Roth A, Sakmann B. 1997. Physiology and anatomy of synaptic connections between thick tufted pyramidal neurones in the developing rat neocortex. J Physiol-Lond 500:409–440.

36. Martini FJ, Guillamón-Vivancos T, Moreno-Juan V, Valdeolmillos M, López-Bendito G. 2021. Spontaneous activity in developing thalamic and cortical sensory networks. Neuron 109:2519– 2534. doi:10.1016/j.neuron.2021.06.026

37. McBride TJ, Rodriguez-Contreras A, Trinh A, Bailey R, DeBello WM. 2008. Learning drives differential clustering of axodendritic contacts in the barn owl auditory system. JNeurosci 28:6960–6973.

38. Meister M, Wong ROL, Baylor DA, Shatz CJ. 1991. Synchronous bursts of action potentials in ganglion cells of the developing mammalian retina. Science 252:939–943.

39. Mel BW, Schiller J, Poirazi P. 2017. Synaptic plasticity in dendrites: complications and coping strategies. *Curr Opin Neurobiol*, Neurobiology of Learning and Plasticity 43:177–186. doi:10.1016/j.conb.2017.03.012

40. Miller M, Peters A. 1981. Maturation of rat visual cortex. II. A combined Golgi-electron microscope study of pyramidal neurons. JComp Neurol 203:555–573.

41. Moyer CE, Zuo Y. 2018. Cortical dendritic spine development and plasticity: insights from in vivo imaging. Curr Opin Neurobiol 53:76–82. doi:10.1016/j.conb.2018.06.002

42. Niculescu D, Michaelsen-Preusse K, Güner Ü, Dorland R van, Wierenga CJ, Lohmann C. 2018. A BDNF-mediated push-pull plasticity mechanism for synaptic clustering. Cell Rep 24:2063–2074. doi:10.1016/j.celrep.2018.07.073

43. Otor Y, Achvat S, Cermak N, Benisty H, Abboud M, Barak O, Schiller Y, Poleg-Polsky A, Schiller J. 2022. Dynamic compartmental computations in tuft dendrites of layer 5 neurons during motor behavior. Science 376:267–275. doi:10.1126/science.abn1421

44. Palmer LM, Shai AS, Reeve JE, Anderson HL, Paulsen O, Larkum ME. 2014. NMDA spikes enhance action potential generation during sensory input. Nat Neurosci 17:383–390. doi:10.1038/nn.3646

45. Pnevmatikakis EA, Giovannucci A. 2017. NoRMCorre: An online algorithm for piecewise rigid motion correction of calcium imaging data. J Neurosci Methods 291:83–94. doi:10.1016/j.jneumeth.2017.07.031

46. Podgorski K, Toth TD, Coleman P, Opushnyev S, Brusco J, Hogg P, Edgcumbe P, Haas K. 2021. Comprehensive imaging of synaptic activity reveals dendritic growth rules that cluster inputs. BioRxiv doi.org/10.1101/2021.02.11.430646. doi:10.1101/2021.02.11.430646

47. Poirazi P, Brannon T, Mel BW. 2003. Pyramidal neuron as two-layer neural network. Neuron 37:989– 999.

48. Poirazi P, Mel BW. 2001. Impact of active dendrites and structural plasticity on the memory capacity of neural tissue. Neuron 29:779–796.

49. Rochefort NL, Garaschuk O, Milos RI, Narushima M, Marandi N, Pichler B, Kovalchuk Y, Konnerth A. 2009. Sparsification of neuronal activity in the visual cortex at eye-opening. Proc Natl Acad Sci 106:15049–15054.

50. Sanes JR, Zipursky SL. 2020. Synaptic Specificity, Recognition Molecules, and Assembly of Neural Circuits. Cell 181:536–556. doi:10.1016/j.cell.2020.04.008

51. Sasaki T, Matsuki N, Ikegaya Y. 2012. Targeted axon-attached recording with fluorescent patch-clamp pipettes in brain slices. Nat Protoc 7:1228–1234. doi:10.1038/nprot.2012.061

52. Scholl B, Thomas CI, Ryan MA, Kamasawa N, Fitzpatrick D. 2021. Cortical response selectivity derives from strength in numbers of synapses. Nature 590:111–114. doi:10.1038/s41586-020-03044-3

53. Siegel F, Heimel JA, Peters J, Lohmann C. 2012. Peripheral and central inputs shape network dynamics in the developing visual cortex in vivo. Curr Biol 22:253–258.

54. Smith SL, Smith IT, Branco T, Häusser M. 2013. Dendritic spikes enhance stimulus selectivity in cortical neurons in vivo. Nature 503:115–120. doi:10.1038/nature12600

55. Svoboda K, Denk W, Kleinfeld D, Tank DW. 1997. In vivo dendritic calcium dynamics in neocortical pyramidal neurons. Nature 385:161–165.

56. Takahashi N, Kitamura K, Matsuo N, Mayford M, Kano M, Matsuki N, Ikegaya Y. 2012. Locally Synchronized Synaptic Inputs. Science 335:353–356. doi: DOI: 10.1126/science.1210362

57. Tran-Van-Minh A, Cazé RD, Abrahamsson T, Cathala L, Gutkin BS, DiGregorio DA. 2015. Contribution of sublinear and supralinear dendritic integration to neuronal computations. Front Cell Neurosci 9:67. doi:10.3389/fncel.2015.00067

58. Voronin LL, Cherubini E. 2004. ‘Deaf, mute and whispering’ silent synapses: their role in synaptic plasticity. J Physiol 557:3–12. doi:10.1113/jphysiol.2003.058966

59. Wildenberg G, Li H, Sampathkumar V, Sorokina A, Kasthuri N. 2023. Isochronic development of cortical synapses in primates and mice. Nat Commun 14:8018. doi:10.1038/s41467-023-43088-3

60. Wilson DE, Whitney DE, Scholl B, Fitzpatrick D. 2016. Orientation selectivity and the functional clustering of synaptic inputs in primary visual cortex. Nat Neurosci 19:1003–1009. doi:10.1038/nn.4323

61. Winfield DA, Brooke RNL, Sloper JJ, Powell TPS. 1981. A combined golgi-electron microscopic study of the synapses made by the proximal axon and recurrent collaterals of a pyramidal cell in the somatic sensory cortex of the monkey. Neuroscience 6:1217–1230. doi:10.1016/0306-4522(81)90183-4

62. Winnubst J, Cheyne JE, Niculescu D, Lohmann C. 2015. Spontaneous activity drives local synaptic plasticity in vivo. Neuron 87:399–410. doi:10.1016/j.neuron.2015.06.029

63. Xu NL, Harnett MT, Williams SR, Huber D, O’Connor DH, Svoboda K, Magee JC. 2012. Nonlinear dendritic integration of sensory and motor input during an active sensing task. Nature 492:247–251.

